# Phenotyping the preterm brain: characterising individual deviations from normative volumetric development in two large infant cohorts

**DOI:** 10.1101/2020.08.05.228700

**Authors:** Ralica Dimitrova, Sophie Arulkumaran, Olivia Carney, Andrew Chew, Shona Falconer, Judit Ciarrusta, Thomas Wolfers, Dafnis Batalle, Lucilio Cordero-Grande, Anthony N. Price, Rui PAG Teixeira, Emer Hughes, Alexia Egloff, Jana Hutter, Antonios Makropoulos, Emma C. Robinson, Andreas Schuh, Katy Vecchiato, Johannes K. Steinweg, Russell Macleod, Andre F. Marquand, Grainne McAlonan, Mary A. Rutherford, Serena J. Counsell, Stephen M. Smith, Daniel Rueckert, Joseph V. Hajnal, Jonathan O’Muircheartaigh, A. David Edwards

## Abstract

The diverse cerebral consequences of preterm birth create significant challenges for understanding pathogenesis or predicting later outcome. Instead of focusing on describing effects common to the group, comparing individual infants against robust normative data offers a powerful alternative to study brain maturation. Here we used Gaussian process regression to create normative curves characterising brain volumetric development in 274 term-born infants, modelling for age at scan and sex. We then compared 89 preterm infants scanned at term-equivalent age to these normative charts, relating *individual* deviations from typical volumetric development to perinatal risk factors and later neurocognitive scores. To test generalisability, we used a second independent dataset comprising of 253 preterm infants scanned using different acquisition parameters and scanner. We describe rapid, non-uniform brain growth during the neonatal period. In *both* preterm cohorts, cerebral atypicalities were widespread, often multiple, and varied highly between individuals. Deviations from normative development were associated with respiratory support, nutrition, birth weight, and later neurocognition, demonstrating their clinical relevance. Group-level understanding of the preterm brain disguise a large degree of individual differences. We provide a method and normative dataset that offer a more precise characterisation of the cerebral consequences of preterm birth by profiling the *individual* neonatal brain.

## Introduction

Preterm birth (or birth before 37 weeks gestational age, GA) affects approximately 10% of pregnancies worldwide (Chawanpaiboon et al. 2019), and is a significant risk predisposing to atypical brain development and lifelong cognitive difficulties including a higher incidence of neurodevelopmental and psychiatric disorders (Nosarti et al. 2012; Agrawal et al. 2018; Thompson et al. 2020). Although early brain correlates of preterm birth have been identified at a group level (Volpe 2019), this vulnerable population is highly heterogeneous, with individuals following diverse clinical and neurocognitive trajectories (Sled and Nossin-Manor 2013; Dimitrova et al. 2020). Indeed, the assumption that prematurity has a homogenous effect on brain development might help account for the relatively poor predictive power of neonatal MRI for later outcome (de Bruïne et al. 2011; Edwards et al. 2018). To better understand brain development, provide accurate prognosis of later functionality, and study the effect of clinical risks and interventions, it is important to provide an *individualised* assessment of cerebral maturation (O’Muircheartaigh et al. 2020). Comparing individuals against robust normative data avoids the requirement to define quasi-homogenous groups in a search for effects common to the group, and offers a powerful alternative to investigate brain development with high sensitivity to pathology at an *individual* infant level (Towgood et al. 2009; Holland et al. 2014; O’Muircheartaigh et al. 2020).

In this study, we used Gaussian process regression (GPR) to create normative charts of typical volumetric development using a large sample of healthy term-born infants scanned cross-sectionally within the first month of life. Analogous to the widely employed paediatric height and weight growth charts, this technique allows the local imaging features of individuals to be referred to typical variation while simultaneously accounting for variables such as age and sex (Marquand et al. 2016, 2019). Having established normative values for brain growth of 14 brain regions, we aimed to (i) quantify deviations from typical development in *individual* preterm infants, (ii) investigate the heterogeneity of these deviations, and (iii) examine the association between individual deviations, perinatal clinical factors and later neurocognitive abilities. To test generalisability, we used a second large independent preterm dataset acquired on a different MR scanner using different imaging parameters.

## Materials and methods

### Participants

This study utilised data from two cohorts. 363 (89 preterm) infants recruited for the developing Human Connectome Project (dHCP; http://developingconnectome.org/) were scanned at term-equivalent age (TEA, 37-45 weeks postmenstrual age, PMA) during natural unsedated sleep at the Evelina London Children’s Hospital between 2015 and 2019. The second cohort comprised of further 253 preterm infants born before 33 weeks GA that underwent MRI between 37 and 45 weeks PMA at the neonatal intensive care unit (NICU) in Hammersmith Hospital between 2010 and 2013 for the Evaluation of Preterm Imaging (EPrime) study. Detailed description of these studies and the scanning procedure used has been previously reported (dHCP (Hughes et al. 2017), EPrime (Edwards et al. 2018)). All MRI images were examined by a neonatal neuroradiologist. Exclusion criteria for term-born infants are described in *Supplementary Methods*. There were no exclusion criteria for the preterm infants, except for major congenital malformations and data included infants from nonsingleton pregnancies. Both studies were approved by the National Research Ethics Committee (dHCP, REC: 14/Lo/1169; EPrime, REC: 09/H0707/98). Informed written consent was given by parents prior to scanning.

### MRI acquisition and preprocessing

MRI data for the dHCP were collected on a Philips Achieva 3T (Philips Medical Systems, Best, The Netherlands) using a dedicated 32-channel neonatal head coil (Hughes et al. 2017). T_2_-weighted scans were acquired with TR/TE of 12s/156ms, SENSE=2.11/2.58 (axial/sagittal), 0.8×0.8mm in-plane resolution, 1.6mm slice thickness (0.8mm overlap). Images were motion corrected and super-resolution reconstructed resulting in 0.5mm isotropic resolution (Makropoulos et al. 2018). MRI data collected for EPrime were acquired on a Philips 3T system using an eight-channel phased array head coil. T_2_-weighted turbo spin echo was acquired with TR/TE of 8670/160ms, in plane resolution 0.86×0.86mm, 2mm slice thickness (1mm overlap).

Both datasets were preprocessed using the dHCP structural pipeline (Makropoulos et al. 2018). In brief, motion-corrected, reconstructed T_2_-weighted images were corrected for bias-field inhomogeneities, brain extracted and segmented. Tissue labels included cerebrospinal fluid (CSF), white matter (WM), cortical grey matter (cGM), deep grey matter (dGM), ventricles (including the cavum, a transient fluid-filled cavity located in the midline of the brain between the left and right anterior horns of the lateral ventricles, which if present, usually disappears during the neonatal period), cerebellum, brainstem, hippocampus and amygdala. dGM was further parcellated into left/right caudate, lentiform and thalamus. Total tissue volume (TTV) incorporated all brain GM and WM volumes; total brain volume (TBV) included TTV and ventricles; intracranial volume (ICV) included TBV and CSF (Table 1.). Given the high correlation between TTV and TBV (ρ=0.98), we reported only TTV. Due to their size and lower tissue contrast in the neonatal brain, the amygdala and the hippocampus are prone to segmentation errors and higher partial voluming, especially in the EPrime dataset, where the image resolution was lower (1mm) compared to dHCP (0.5mm). Therefore, these structures were excluded from the present analyses. The quality of the preprocessing was visually evaluated using a scoring system detailed elsewhere (Makropoulos et al. 2018) to ensure no images severely affected by motion or with poor segmentation were included (*Supplementary Methods & Supplementary Figure 1*).

**Table 1.** Brain regions of interest, the structures they include and what global brain measures they are taken as a proportion from when calculating relative brain volumes.

| Brain regions | Brain structures included | Relative volume (% from) |
| --- | --- | --- |
| cortical Grey Matter (cGM) | - | TTV |
| White Matter (WM) | - | TTV |
| Cerebellum | - | TTV |
| Brainstem | - | TTV |
| Cerebrospinal fluid (CSF) | - | ICV |
| Ventricles | Lateral ventricles + cavum | TBV |
| Caudate (left/right) | - | TTV |
| Lentiform (left/right) | Pallidum + Putamen | TTV |
| Thalamus (left/right) | - | TTV |
| Total tissue volume (TTV) | All brain GM + WM tissue | - |
| Total brain volume (TBV) | All brain GM + WM tissue + ventricles | - |
| Intracranial volume (ICV) | All brain GM + WM tissue + ventricles + CSF | - |

We estimated regional volumes in absolute (cm^3^) and relative (%) values. Relative volumes were calculated as the proportion of each tissue volume from TTV, the ventricles from TBV and CSF from ICV (Table 1). To capture the effect of preterm birth, we used relative volumes to (i) ensure results are not driven by extreme individual differences in non-brain intracranial volume, often seen in preterm infants, (ii) partially alleviate differences in data acquisition.

### Modelling volumetric development using Gaussian Process Regression

To characterise neonatal volumetric development, we used GPR, a Bayesian non-parametric regression, implemented in GPy (https://sheffieldml.github.io/GPy/). GPR simultaneously provides point estimates and measures of predictive confidence for every observation representing the distance of each individual observation from the normative mean at that point on the ‘curve’ accounting for modelled covariates (Marquand et al. 2016).

We first trained a GPR model to describe typical development in the term-born dataset (274 infants) using PMA at scan and sex to predict 14 brain structures separately. Regions included ICV, TTV, cGM, WM, cerebellum, brainstem, CSF, ventricles and left/right caudate, lentiform and thalamus. Model accuracy was tested under 5-fold cross-validation, with each fold stratified to cover the whole PMA range (37-45 weeks). The relationship between the volume outputs and model predictors was estimated with a sum of radial basis function, linear and white noise covariance kernels. Model hyperparameters were optimised using log marginal likelihood. Prediction performance was evaluated using the mean absolute error (MAE) between the predicted and the observed value derived from the 5-fold cross-validation.

To assess the effects of preterm birth, we retrained the model on the entire term-born dataset and applied the model to 89 dHCP preterm infants scanned at TEA. To assess generalisability, we applied the same model to 253 preterm infants from the EPrime study. A Z-score was derived for every infant by estimating the difference between the model prediction and the observed value normalised by the model uncertainty (the square root of the predicted variance). To quantify extreme deviations, prior to analyses, we chose a threshold of |Z|>2.6 (corresponding to p<0.005) following the convention adopted in previous GPR analyses modelling adult brain development (Wolfers et al. 2019). We examined the proportion of infants with volumes lying more than 2.6 standard deviations (sd) above/below the model mean (indicating the top/bottom 0.5% of the typical group values, hereafter described as extreme positive or negative deviations, respectively).

To quantify the effect of image spatial resolution differences between the dHCP and EPrime, we first downsampled the dHCP data to 1mm isotropic resolution using FSL flirt (-applyisofxm, spline interpolation), reran the tissue segmentation on the new dHCP resolution data and trained the GPR model. We examined the difference in (i) model means and (ii) the number of EPrime infants who deviated significantly from the predicted model means.

### Deviations from normative development, perinatal risks and later neurocognition

We tested the association between deviations from normative development (in relative volumes) and recognised perinatal clinical risks (Boardman and Counsell 2019), including GA at birth, birth weight Z-score, total days receiving mechanical ventilation, continuous positive airway pressure (CPAP) and total parenteral nutrition (TPN, available only for EPrime). Birth weight Z-scores were calculated using the population data from the uk90 growth charts implemented in sitar R package (Cole et al. 2010). Oxygen/respiratory support and nutrition were administered at the NICU. These data were obtained from electronic hospital records and days were counted if the infant spent any part of the day on ventilation, CPAP or TPN, with higher number of days indicative of poorer health. Bayley III Scales of Infant Development (BSID-III) (Bayley 2006) assessment was carried out by trained developmental paediatricians/psychologists at 18 months for the dHCP and at 20 months for EPrime (corrected age). We used the composite scores for motor, cognitive and language development (mean(sd)=100(15)). Associations were examined using Spearman rho (r) or Mann-Whitney U test combined with Cliff’s delta (d, ranging from −1 to 1), under Bonferroni-Holm multiple comparison corrections.

### Data availability

Normative term-born data and the GPR code used in this study are freely available on GitHub (https://github.com/ralidimitrova). All imaging data collected for the dHCP will be publicly available in early 2021 at http://developingconnectome.org/.

## Results

The perinatal, demographic and neurocognitive characteristics are presented in Table 2. EPrime infants were born earlier (p<0.05, d=0.28) and had lower birth weight (p<0.05, d=0.19) compared to dHCP preterm infants. On average, they had poorer motor (p<0.05, d=0.22) skills at follow-up (language p=0.1; cognition p=0.12). There were no differences in days on CPAP (p=0.57), but dHCP preterm infants required mechanical ventilation for longer (p<0.05, d=0.16). The two preterm cohorts did not differ in punctate WM lesions (PWMLs, p=0.09) or cerebellar haemorrhages (p=0.82) incidence, nor in proportion of infants with intrauterine growth restriction (IUGR, p=0.15); yet haemorrhagic parenchymal infarction (HPI) and periventricular leukomalacia (PVL) were observed only in the EPrime cohort (Table 2.).

**Table 2.**
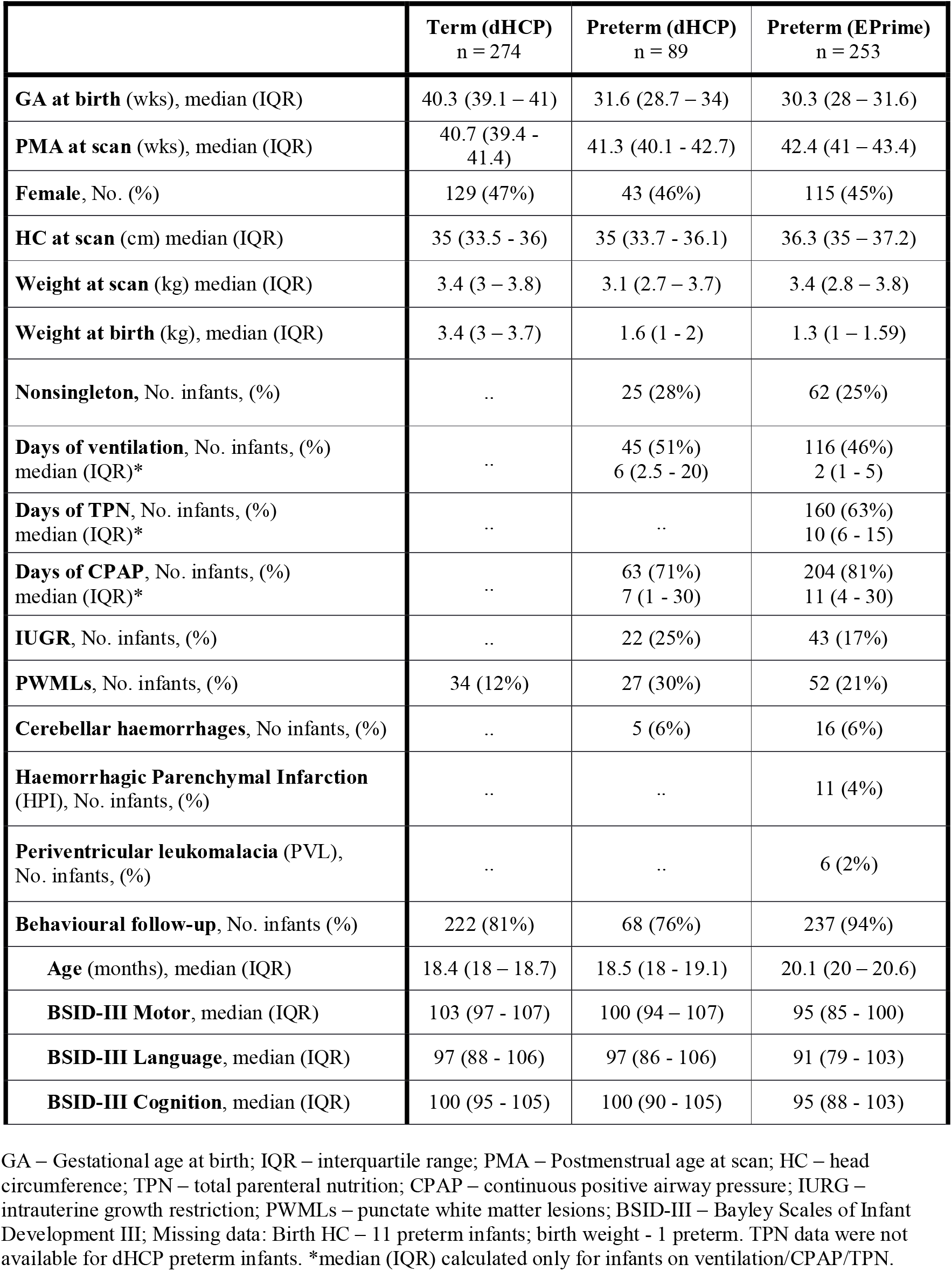
Perinatal, demographic and neurocognitive characteristics of the study sample.

|  | **Term (dHCP)**<br>n = 274 | **Preterm (dHCP)**<br>n = 89 | **Preterm (EPrime)**<br>n = 253 |
| --- | --- | --- | --- |
| **GA at birth** (wks), median (IQR) | 40.3 (39.1 – 41) | 31.6 (28.7 – 34) | 30.3 (28 – 31.6) |
| **PMA at scan** (wks), median (IQR) | 40.7 (39.4 - 41.4) | 41.3 (40.1 - 42.7) | 42.4 (41 – 43.4) |
| **Female**, No. (%) | 129 (47%) | 43 (46%) | 115 (45%) |
| **HC at scan** (cm) median (IQR) | 35 (33.5 - 36) | 35 (33.7 - 36.1) | 36.3 (35 – 37.2) |
| **Weight at scan** (kg) median (IQR) | 3.4 (3 – 3.8) | 3.1 (2.7 – 3.7) | 3.4 (2.8 – 3.8) |
| **Weight at birth** (kg), median (IQR) | 3.4 (3 – 3.7) | 1.6 (1 - 2) | 1.3 (1 – 1.59) |
| **Nonsingleton**, No. infants, (%) | .. | 25 (28%) | 62 (25%) |
| **Days of ventilation**, No. infants, (%)<br>median (IQR)* | .. | 45 (51%)<br>6 (2.5 - 20) | 116 (46%)<br>2 (1 - 5) |
| **Days of TPN**, No. infants, (%)<br>median (IQR)* | .. | .. | 160 (63%)<br>10 (6 - 15) |
| **Days of CPAP**, No. infants, (%)<br>median (IQR)* | .. | 63 (71%)<br>7 (1 - 30) | 204 (81%)<br>11 (4 - 30) |
| **IUGR**, No. infants, (%) | .. | 22 (25%) | 43 (17%) |
| **PWMLs**, No. infants, (%) | 34 (12%) | 27 (30%) | 52 (21%) |
| **Cerebellar haemorrhages**, No infants, (%) | .. | 5 (6%) | 16 (6%) |
| **Haemorrhagic Parenchymal Infarction (HPI)**, No. infants, (%) | .. | .. | 11 (4%) |
| **Periventricular leukomalacia (PVL)**,<br>No. infants, (%) | .. | .. | 6 (2%) |
| **Behavioural follow-up**, No. infants (%) | 222 (81%) | 68 (76%) | 237 (94%) |
| **Age** (months), median (IQR) | 18.4 (18 – 18.7) | 18.5 (18 - 19.1) | 20.1 (20 – 20.6) |
| **BSID-III Motor**, median (IQR) | 103 (97 - 107) | 100 (94 – 107) | 95 (85 - 100) |
| **BSID-III Language**, median (IQR) | 97 (88 - 106) | 97 (86 - 106) | 91 (79 - 103) |
| **BSID-III Cognition**, median (IQR) | 100 (95 - 105) | 100 (90 - 105) | 95 (88 - 103) |

GA – Gestational age at birth; IQR – interquartile range; PMA – Postmenstrual age at scan; HC – head circumference; TPN – total parenteral nutrition; CPAP – continuous positive airway pressure; IURG – intrauterine growth restriction; PWMLs – punctate white matter lesions; BSID-III – Bayley Scales of Infant Development III; Missing data: Birth HC – 11 preterm infants; birth weight - 1 preterm. TPN data were not available for dHCP preterm infants. \*median (IQR) calculated only for infants on ventilation/CPAP/TPN.

### Typical volumetric development in term-born infants during the neonatal period

We found an increase in all absolute volumes except the ventricles, where no change was detected (Fig. 1A, *Supplementary Figure 2*). The increase was greatest in cGM (10.4% per week, pw) and cerebellum (9.9% pw) compared to ICV (6.1 % pw), TTV (6% pw) and CSF (7% pw). Subcortical structures increased between 4 and 6% pw (caudate L:4.1%, R:4.3%; lentiform L:6.6%, R:5.3%; thalamus L:4.1%, R:4.7% pw) with smaller increases in brainstem (3.9% pw) and WM (2% pw).

**Figure 1.**
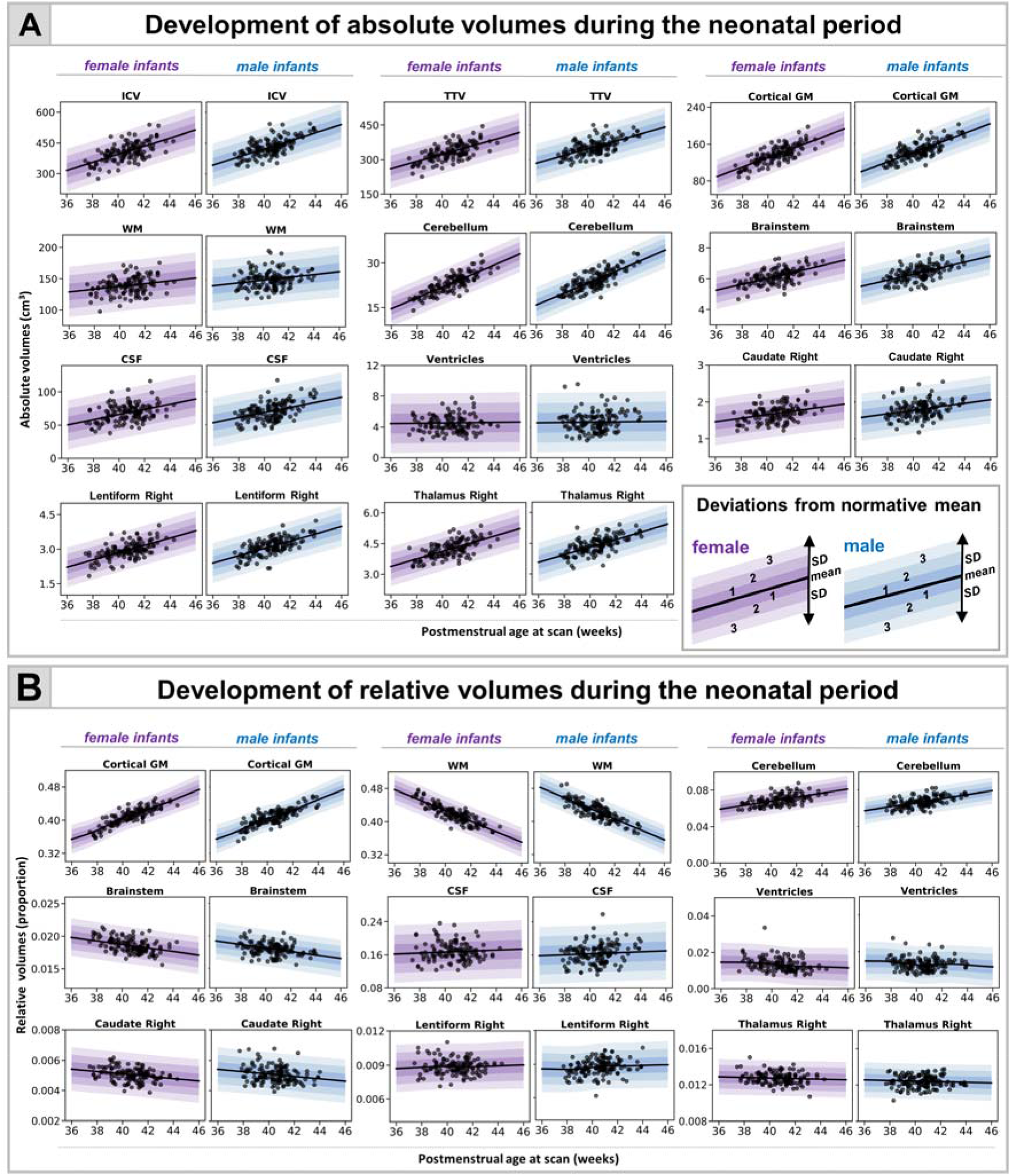
Normative modelling of volumetric development during the neonatal period. The model means for both female and male term infants are shown in purple and blue respectively together with ±1, ±2 and ±3 standard deviations from the model means for absolute (A) and relative (B) volumes (tissue volumes represented as a proportion from TTV, ventricles from TBV and CSF from ICV). Normative charts are shown only for right dGM structures (left structures are shown in *Supplementary Figure 2*).

The greatest changes in relative volumes were observed in cGM and WM (Fig. 1B, *Supplementary Figure 2*). cGM represented 36% of TTV at 37 weeks PMA, and increased to 44% at 44 weeks PMA, while the relative WM volume decreased from 48% to 38% of TTV. The relative cerebellar volume increased from 6% to 7%. There was an increase in relative lentiform volume, a subtle decrease in caudate and no change in thalamus. We observed a slight increase in CSF proportion of ICV and a steady decrease in the proportion that ventricles contributed to TBV. MAE for all models is shown in *Supplementary Table 1*.

### Image resolution and volumetric development

Overall, the majority of observations in both dHCP and EPrime preterm samples fit within 2.6 sd of the term-born model, indicating good agreement between the two studies (Fig. 2A). Differences were most profound in fluid-filled structures, likely attributable to partial voluming of high T_2_-signal CSF. In agreement, when compared to the models built using the original dHCP resolution of 0.5mm, the matched 1mm resolution models showed a mean shift (increase) for the CSF and ventricular volumes (Fig. 2B; *Supplementary Figure 3 & 4*). As a result, when using the lower resolution normative charts, the proportion of extreme positive deviations in EPrime decrease from 53% to 29% for CSF and from 44% to 32% for ventricles (Fig. 2C). All infants who showed extreme deviations in the matched 1mm resolution showed extreme deviations in the original 0.5mm resolution. Changes in the proportion of extreme deviations associated with image resolution for the rest of the structures were more subtle. Unless stated otherwise, data are presented for the 0.5mm resolution models. The overall proportion of extreme deviations in the term-born sample was very low in all brain structures, with no more than 2% of the sample with Z-scores > |2.6| in the original 0.5mm resolution.

**Figure 2.**
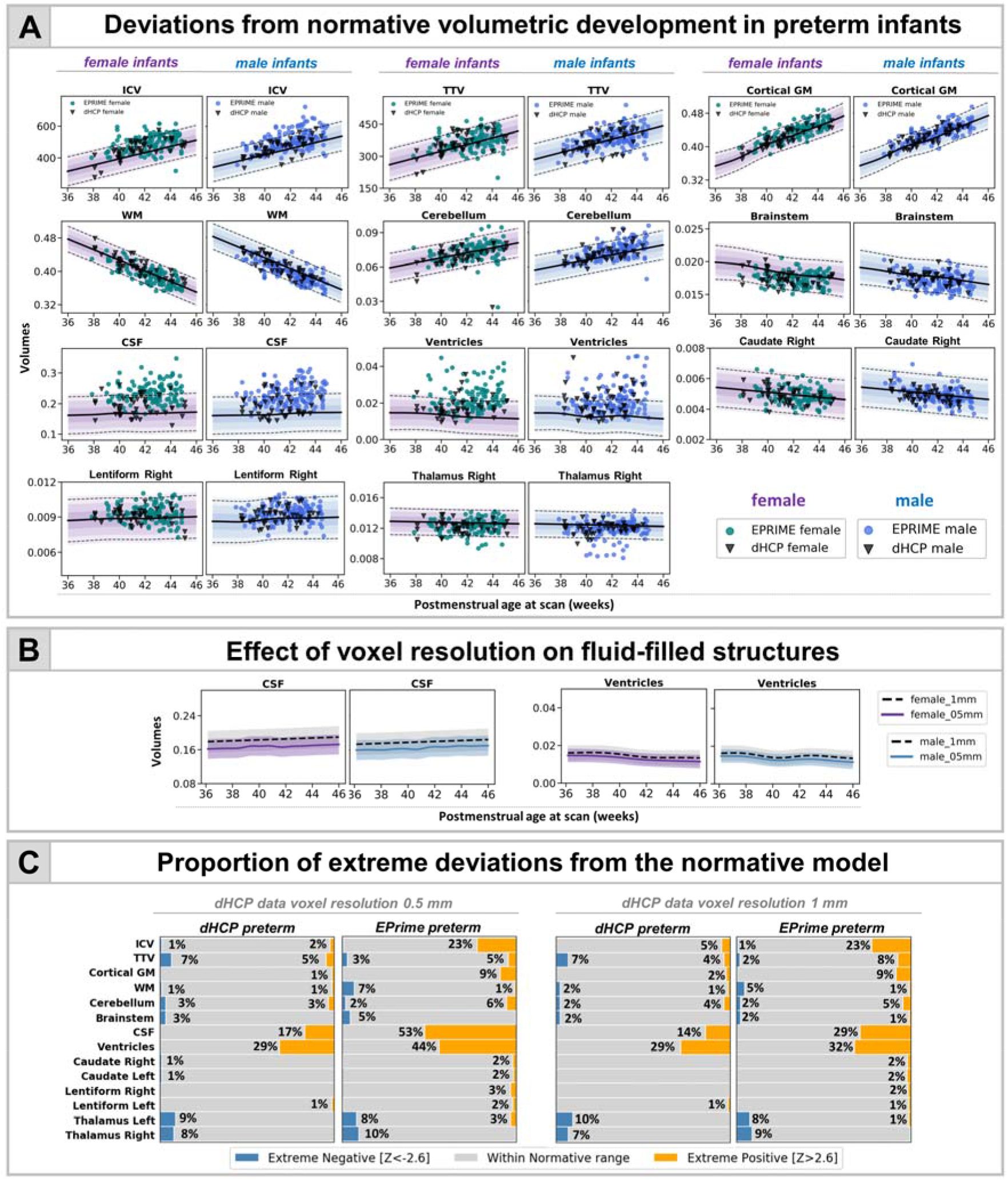
Characterising the effects of preterm birth on the developing brain. (A) Deviations from normative volumetric development in preterm infants. Observations for individual preterm infants from both dHCP and EPrime cohorts are shown with model means for both female and male term-born infants together with ±1, ±2 and ±3 standard deviations. Intracranial brain volume (ICV), total brain volume (TBV) and total tissue volume (TTV) are in cm^3^; cGM, WM, cerebellum, brainstem and subcortical structures shown as a proportion of TTV; CSF as a proportion of ICV and ventricles as a proportion of TTV. Horizontal --- lines show Z > |2.6|, the threshold used to define extreme deviations. The normative curves for the ventricles show data within 10 sd from the mean, full range is shown in Fig. 4 and discussed below. (B) Mean differences in fluid filled structures between GPR models build using 0.5mm and 1mm dHCP imaging resolution. (C) Proportion (%) of extreme deviations from the normative model in preterm infants. Extreme negative deviations (Z < −2.6) are depicted in blue, while extreme positive deviations (Z > 2.6) are shown in orange.

### Infants with reduced total tissue volumes suffered more extreme prematurity

Six dHCP preterm infants (7%) and eight EPrime (3%) infants showed extreme negative deviations in TTV (Fig. 2C), all of which (except one EPrime infant) were born at GA<30 weeks and weighed less than 1 kg at birth. Significantly reduced TTV was accompanied by enlarged CSF and ventricles (dHCP: 3/6; EPrime: 7/8). Four infants (dHCP: 1/6; EPrime 3/8) also had associated reduction in cerebellar volumes. A further three preterm infants (dHCP: 1/6; EPrime 2/8) had associated reduction in the thalami, bilaterally. All infants required oxygen support after birth. Infants who had TTV 2.6 sd above the model mean (dHCP: 4 (5%); EPRIME: 18 (7%)), were born GA>30 weeks and had no incidental findings, other than PWMLs, and short need for oxygen support and TPN after birth.

### Infants with reduced thalamic volume also had PWMLs

In the dHCP preterm sample, all eight infants with extreme negative thalamic deviations had PWMLs, seven of eight had multiple lesions. Four out of these seven infants had lesions involving the corticospinal tract (Fig. 3). Seven out of the eight infants were on CPAP, but none of them for a long period of time (five infants <4 days; one infant 11 days; one infant 18 days) and all seven did not require ventilation. None of these infants had a birthweight of less than 1kg. In EPrime 17 infants had bilateral reduced thalamic volume and 10 unilateral extreme deviations (with structure in the other hemisphere close to but not reaching Z<-2.6). 78% of these infants had PWMLs compared to 16% incidence in the rest of the sample. Overall, across the whole cohort, infants with PWMLs had significantly reduced left (d=0.56) and right (d=0.53) thalamic volumes, compared to infants without (both p<0.05). In EPrime, infants with reduced thalamic volumes, often had CSF or ventricular volumes significantly bigger than the normative values for their age/sex. In five infants, this was associated with PVL and in a further two, with HPI.

**Figure 3.**
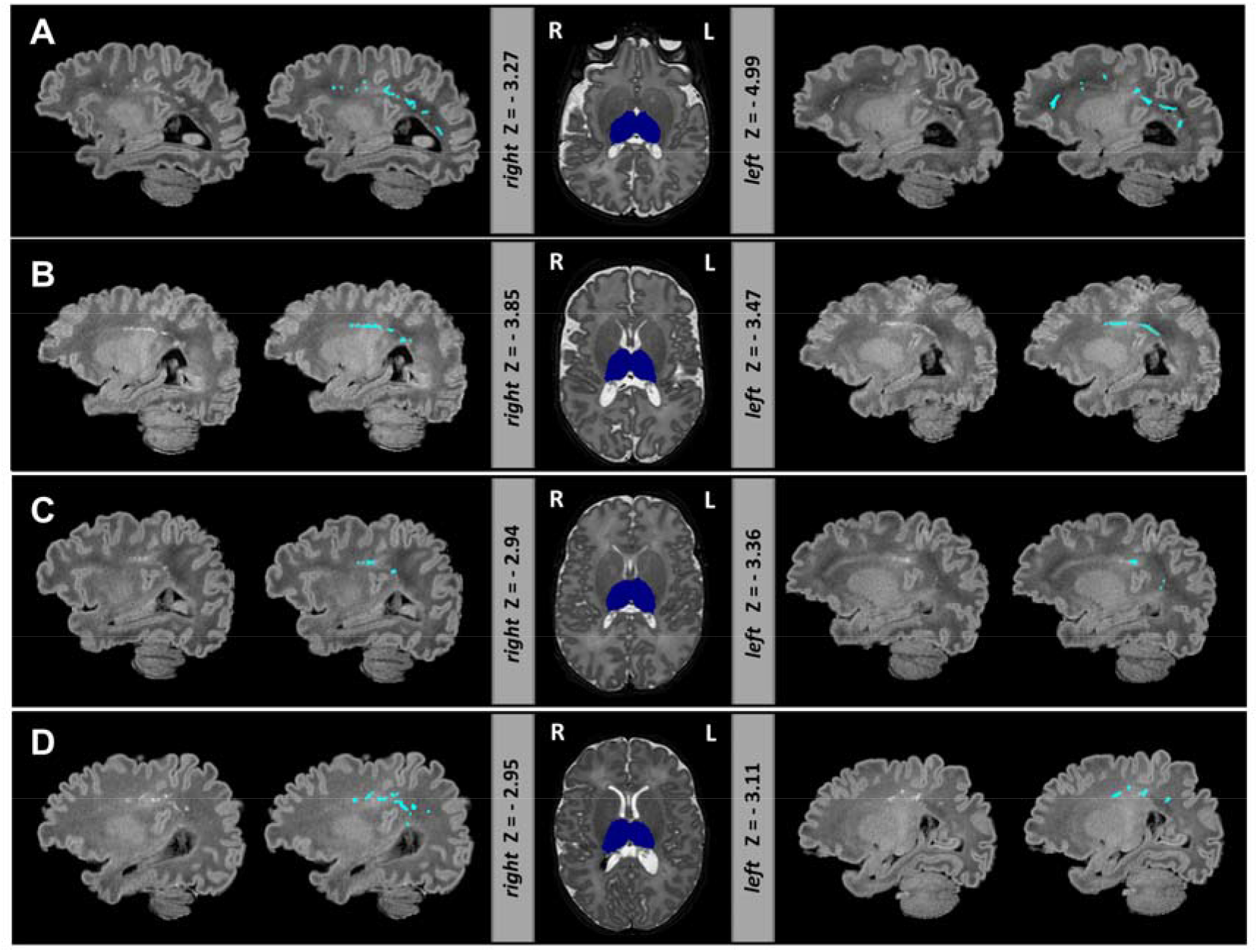
Extreme negative deviations in thalamic volume were often accompanied by punctate WM lesions in the preterm brain. Depicted are four infants (A-D) with bilateral thalamic volumes significantly below the model mean. Thalamic segmentation (dark blue) is overlaid onto the T_2_-weighted images. T1-weighted images are shown with and without the manual outlined punctate WM lesions (light blue). Note T1-weighted images were not used in the preprocessing but are shown here due to better contrast for detecting PWMLs.

### Atypical ventricular development in preterm infants: frequent but highly heterogeneous

Widening of the fluid-filled structures was the most frequently observed deviation from normative development in both cohorts. In the dHCP 29% and 17% of the preterm infants showed extreme deviations in ventricular and CSF volumes, respectively. This number was higher in EPrime where increased ventricles and CSF were seen in 44% and 53% of infants with the original 0.5mm dHCP resolution and in 29% and 32% with the downsampled 1mm resolution. Figure 4 shows the most extreme cases where infants’ ventricles were 10 sd above the model mean. These extreme deviations in ventricular volume were associated with overt focal brain injuries including haemorrhagic parenchymal infarction (infants 1,2,4,6) and periventricular leukomalacia (infants 5,7). In all of these infants we also observed significant negative deviations in TTV or thalamus and increased CSF. These infants performed poorly at follow-up (Fig. 4).

**Figure 4.**
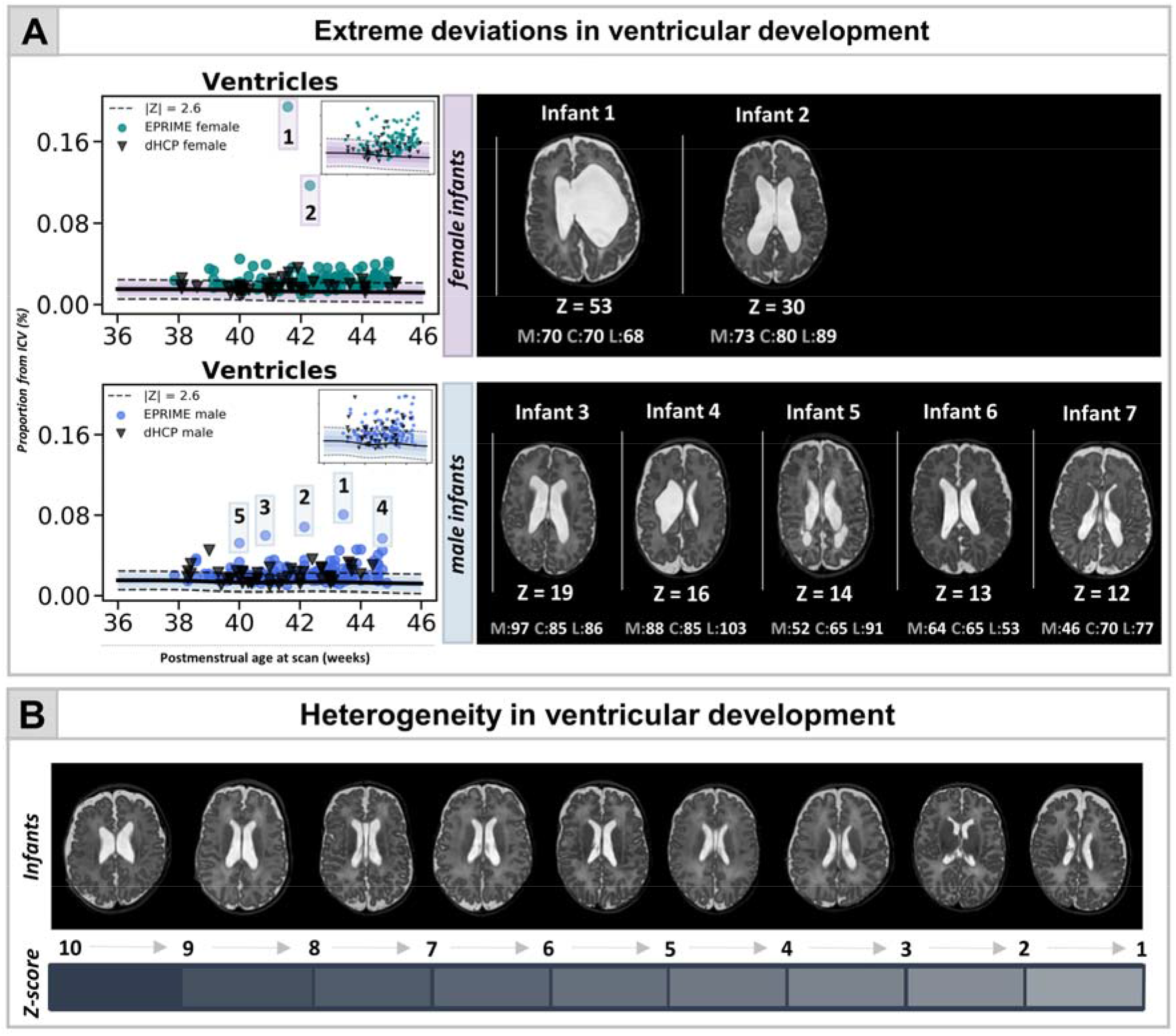
Capturing heterogeneity and extreme deviations in ventricular development in the preterm brain at term-equivalent age. (A) Normative curves are shown for both female and male infants (in upper right corner – curves excluding the outliers, also shown in Fig. 2A). The figure also depicts the T_2_-weighted images for infants with ventricular volume lying 10 sd above the mean, separate for females (top) and males (bottom), together with their neurocognitive scores (M – motor, C – cognitive, L – language). Ventricular development in EPrime preterm infants is highly heterogeneous both in shape and size as illustrated in (B) showing ventricular volumes of various Z-scores.

### Association between perinatal risks and deviations from normative development

In the dHCP cohort, decreased GA at birth related to reduced TTV (ρ=0.45) and increased proportion of CSF (ρ=−0.44), while in EPrime to reduced TTV (ρ=0.27) and increased relative ventricular volumes (ρ=−0.26) (all p_corr_<0.05) (Fig. 5; *Supplementary Table 2*). In both samples, greater birth weight Z-score was related to bigger ICV (dHCP: ρ=0.41, EPrime: ρ=0.36) and TTV (ρ=0.40, ρ=0.37) at TEA and in EPrime alone, to reduced relative brainstem (ρ=−0.26) and bilateral thalamic volumes (right: ρ=−0.28, left: ρ=−0.25) (all p_corr_<0.05; *Supplementary Table 3*).

**Figure 5.**
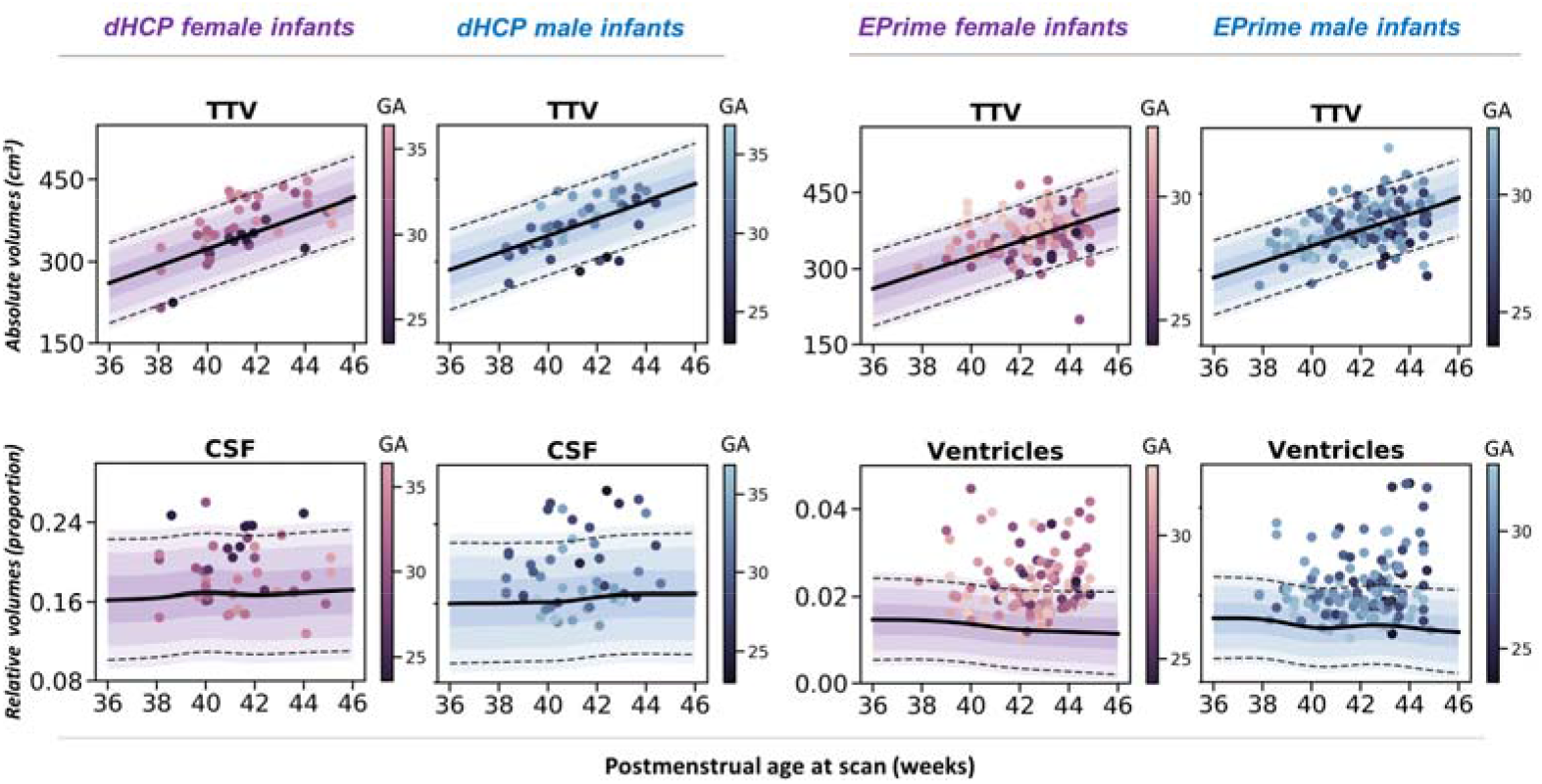
Association between degree of prematurity and deviations from normative brain development. In the dHCP preterm sample, increased degree of prematurity (lower GA at birth) was related to reduced TTV and increased CSF. In the EPrime sample, increased degree of prematurity was associated with reduced TTV and increased ventricular volume. Individual preterm observations are plotted against the normative model mean for female (purple) and male (blue) term infants. The plots also show ±1, ±2 and ±3 standard deviations from the normative means together with --- lines indicating Z > |2.6|, the threshold used to define extreme deviations. Ventricular data are shown only for infants with volume ±10 standard deviations from the model mean.

In the dHCP preterm sample, longer requirement for CPAP related to smaller TTV (ρ=−0.47), and ICV (ρ= −0.37) as well as to increased proportion CSF (ρ=0.39) and ventricles (ρ=0.37). Longer need for mechanical ventilation was associated with reduced TTV (ρ=−0.44) and relative left caudate volume (ρ=−0.34) as well as with increased relative CSF (ρ=0.39) and ventricular volumes (ρ=0.42) (all p_corr_<0.05). Consistent with this, in EPrime, longer requirement for both CPAP and mechanical ventilation were related to reduced TTV (ρ=−0.30, ρ=−0.38, respectively) and increased relative ventricular volume (both at ρ=0.22) (all p_corr_<0.05). Increased number of days requiring TPN were related to reduced TTV (ρ=−0.43, p_corr_<0.05) and increased relative ventricular volume (ρ=0.25, p_corr_<0.05; *Supplementary Table 4*).

### Association between deviations from normative development and neurocognitive outcome

In the dHCP preterm sample, increased ventricular volume was related to poorer motor (ρ=−0.40) and language (ρ=−0.37) scores at 18 months (all p_corr_<0.05). In EPrime, increased CSF (ρ=−0.22) and cGM (ρ=−0.21), and reduced WM (ρ=0.20) were associated with poorer language abilities, while reduced TTV (ρ=0.20), increased CSF (ρ=−0.25) and cGM (ρ=−0.20) were linked to poorer cognitive performance (all p_corr_<0.05). While most of these association were of similar effect size in the dHCP, they did not survive multiple comparison correction (*Supplementary Table 5*).

## Discussion

The diverse cerebral consequences of preterm birth create significant challenges for understanding pathogenesis or predicting later neurocognitive outcomes. Focusing on individuals and their unique cerebral development can offer new insights. In this study, we first characterised normative volumetric development during the neonatal period, to then describe the effect of preterm birth at an *individual* infant level. We showed deviations from the normative curves consistent with previous studies but with marked variability among individuals. These individual deviations were associated with perinatal risks and later neurocognition.

We previously demonstrated that GPR could be used to detect subtle WM injury with high sensitivity (O’Muircheartaigh et al. 2020), and to characterise the heterogeneous consequences of preterm birth on the developing brain microstructure (Dimitrova et al. 2020). The present application of GPR to volumetric data offers more straightforward clinical translation. GPR provides normative charts describing typical volumetric development and can detect and quantify atypical maturation in individual infants (Ou et al. 2017). The GPR approach generalised to a cohort of infants with MRI data collected on a different MR scanner with different acquisition parameters. In the future, this method could be integrated into automatic tools that complement radiological decisions regarding infant development (Duerden and Thompson 2020). Our method and normative dataset are freely available for researchers to use for understanding pathogenesis, trialling interventions and defining neurocognitive prognosis for vulnerable preterm infants.

We quantified rapid postnatal brain growth consistent with previous imaging and post-mortem studies describing change in the size, organisation and complexity of the brain during the perinatal period (Huttenlocher and Dabholkar 1997; Hüppi et al. 1998; Knickmeyer et al. 2008; Kersbergen et al. 2016; Makropoulos et al. 2016). This dramatic growth is a sum product of a number of heterochronous developmental processes that take place in the developing brain including synaptogenesis, dendritic arborization and early stages of myelination (Huttenlocher 1990; Huttenlocher and Dabholkar 1997; Travis et al. 2005; Petanjek et al. 2008; Lebenberg et al. 2019). Abrupt preterm extrauterine exposure represents a significant stressor to these events and may lead to widespread deviations from the normative trajectories in any or many of these processes as seen in pathology and pre-clinical models (Elovitz and Mrinalini 2004; Burd et al. 2012; Volpe 2019) associated with atypical trajectory of brain growth and a wide range of neurodevelopmental consequences (Inder et al. 2005; Bora et al. 2014; Ball et al. 2017; Gui et al. 2019). However, these alterations are not a result of loss of intrauterine environment alone but are a product of the cumulative effects of clinical and genetic factors creating individualised circumstances for every infant. GPR applied to a large normative dataset offers a powerful approach to study how preterm birth shapes the brain at an *individual* infant level and offers the means to capture important differences in single infants that may be missed by analysis of the means/medians of quasi-homogenous groups which ‘averages-out’ personal effects.

By quantifying this inter-individual variability, our analysis clarified the relationship between reduced global brain growth and preterm birth. Many but not all studies show group-level differences in TTV between preterm and term-born infants (Boardman et al. 2007). We report a subset of infants in both preterm cohorts that deviated significantly from normative brain volumes. These infants were born very early, very small and had prolonged need for supplemental oxygen. Consistent with this, lower GA at birth, birthweight Z-score, longer requirement for respiratory support and TPN were related to reduced TTV and enlarged CSF/ventricles in both preterm cohorts. Longitudinal studies suggest that these effects are not only evident at TEA but might persist to childhood and later life (Nosarti 2002; Allin et al. 2004; Ment et al. 2009; de Kieviet et al. 2012; El Marroun et al. 2020). Not all extremely preterm infants had TTV deviations significantly below the model mean, which could explain the discrepancies found between previous group analyses studying the association between preterm birth and reduced brain volume. An individualised approach is now possible to address the important question of which protective factors or lack of adverse perinatal risks, lead to typical global brain growth in these at-risk infants.

The period encompassing mid gestation and the last trimester of pregnancy is a critical phase for the development and establishment of the thalamocortical network (Kostović et al. 2014). During this short period, there are dynamic changes in thalamocortical efferent fiber organisation and cortical lamination, including rapid axonal growth and the dissolution of the subplate (Kostovic and Rakic 1990; Vasung et al. 2011; Kostović et al. 2014). This makes the thalamus and connecting WM projections particularly vulnerable to injury as a result of preterm birth (Boardman et al. 2006; Ball et al. 2015) with studies suggesting abnormal development may persist beyond TEA (Lin et al. 2001). We reported a subset of preterm infants with thalamic volumes significantly below the model mean (Z<<2.6). These infants had a high load of PWMLs, and five of the EPrime infants had PVL, supporting previous findings of a close link between thalamic development and WM abnormalities, including a previous group analysis of the EPrime dataset (Boardman et al. 2006; Pierson et al. 2007; Ligam et al. 2009; Volpe 2009; Ball et al. 2015; Wisnowski et al. 2015; Tusor et al. 2017). The exact mechanisms that underlie reduced thalamic growth, possibly including neuronal loss and/or atypical developmental trajectory triggered by preterm extrauterine exposure, however remain elusive (Volpe 2009).

Compared to the dHCP preterm cohort, the EPrime study comprised extremely preterm infants, that were sicker during clinical care, had overall poorer motor outcomes, and were scanned using different acquisition parameters. These factors in combination likely underlie some of the differences in associations between extreme deviations and later neurocognitive scores observed between the two datasets. The lower spatial resolution in EPrime in particular, contributed to the mean shift (increase) in CSF and ventricular volumes observed in the EPrime. With all this in mind, it was reassuring that deviations in brain development and their association with perinatal risks found in the dHCP broadly replicated in EPrime, indicating good generalisation of the model to independent data collected on a different MRI scanner. We chose to use volumetric measures that are easy to calculate in research studies or routine clinical examinations. This could offer a direct clinical application, though given the regional heterochrony of early brain development (Lebenberg et al. 2019), future work should focus on more finely-parcellated regions or more sophisticated MRI-derived features, including cortical thickness and surface area. We reported an association between deviations from normative brain development at TEA and behaviour at 18-20 months. An important step for future research is to investigate whether these early brain deviations persist in later life and are predictive of childhood and later neurodevelopment (Boardman et al. 2020; George et al. 2020).

The argument that every brain is different is not novel, and the expectation that the effects of preterm birth are homogeneous and exactly alike in every infant is equally untenable. Individualised methodologies have been successfully applied in other fields (e.g. neuropsychology (Towgood et al. 2009), ageing (Ziegler et al. 2014)) and hold significant promise for the preterm infant. Although a group-mean difference is detectable using the conventional case-control approach, the significant heterogeneity would not be captured and effects of clinical significance to individual infants would be averaged out (Sled and Nossin-Manor 2013). Additionally, visually subtle effects may have prognostic significance when combined with other deviations from normative brain growth, for example reduced thalamic volume, and further analytic power may be gained by including covariates in the GPR model.

In summary, our approach offers a readily interpretable, generalisable and more precise understanding of the cerebral consequences of preterm birth by focusing on the individual rather than the group average atypicality, and in future might improve the predictive power of neuroimaging.

## Acknowledgements

The authors would like to thank all the families who dedicated their time to take part in these studies. The authors declare no conflict of interest.

## References

Agrawal S, Rao SC, Bulsara MK, Patole SK. 2018. Prevalence of Autism Spectrum Disorder in Preterm Infants: A Meta-analysis. Pediatrics. 142:e20180134.

Allin M, Henderson M, Suckling J, Nosarti C, Rushe T, Fearon P, Stewart AL, Bullmore ET, Rifkin L, Murray R. 2004. Effects of very low birthweight on brain structure in adulthood. Dev Med Child Neurol. 46:46–53.

Ball G, Aljabar P, Nongena P, Kennea N, Gonzalez-Cinca N, Falconer S, Chew ATM, Harper N, Wurie J, Rutherford MA, Counsell SJ, Edwards AD. 2017. Multimodal image analysis of clinical influences on preterm brain development. Ann Neurol. 82:233–246.

Ball G, Pazderova L, Chew A, Tusor N, Merchant N, Arichi T, Allsop JM, Cowan FM, Edwards AD, Counsell SJ. 2015. Thalamocortical connectivity predicts cognition in children born preterm. Cereb cortex. 25:4310–4318.

Bayley N. 2006. Bayley Scales of Infant and Toddler Development. San Antonio: TX The Psychological Corporation.

Boardman JP, Counsell SJ. 2019. Invited Review: Factors associated with atypical brain development in preterm infants: insights from magnetic resonance imaging. Neuropathol Appl Neurobiol. 44.

Boardman JP, Counsell SJ, Rueckert D, Hajnal J V., Bhatia KK, Srinivasan L, Kapellou O, Aljabar P, Dyet LE, Rutherford MA, Allsop JM, Edwards AD. 2007. Early growth in brain volume is preserved in the majority of preterm infants. Ann Neurol. 62: 185–192.

Boardman JP, Counsell SJ, Rueckert D, Kapellou O, Bhatia KK, Aljabar P, Hajnal J, Allsop JM, Rutherford MA, Edwardsa AD. 2006. Abnormal deep grey matter development following preterm birth detected using deformation-based morphometry. Neuroimage. 32:70–78.

Boardman JP, Hall J, Thrippleton MJ, Reynolds RM, Bogaert D, Davidson DJ, Schwarze J, Drake AJ, Chandran S, Bastin ME, Fletcher-Watson S. 2020. Impact of preterm birth on brain development and long-term outcome: protocol for a cohort study in Scotland. BMJ Open. 10:e035854.

Bora S, Pritchard VE, Chen Z, Inder TE, Woodward LJ. 2014. Neonatal cerebral morphometry and later risk of persistent inattention/hyperactivity in children born very preterm. J Child Psychol Psychiatry Allied Discip. 55:828–838.

Burd I, Balakrishnan B, Kannan S. 2012. Models of Fetal Brain Injury, Intrauterine Inflammation, and Preterm Birth. Am J Reprod Immunol. 67:287–294.

Chawanpaiboon S, Vogel JP, Moller AB, Lumbiganon P, Petzold M, Hogan D, Landoulsi S, Jampathong N, Kongwattanakul K, Laopaiboon M, Lewis C, Rattanakanokchai S, Teng DN, Thinkhamrop J, Watananirun K, Zhang J, Zhou W, Gülmezoglu AM. 2019. Global, regional, and national estimates of levels of preterm birth in 2014: a systematic review and modelling analysis. Lancet Glob Heal. 7:e37–e46.

Cole TJ, Donaldson MDC, Ben-shlomo Y. 2010. SITAR-a useful instrument for growth curve analysis. Int J Epidemiol. 39:1558–1566.

de Bruïne FT, van den Berg-Huysmans AA, Leijser LM, Rijken M, Steggerda SJ, van der Grond J, van Wezel-Meijler G. 2011. Clinical implications of MR imaging findings in the white matter in very preterm infants: a 2-year follow-up study. Radiology. 261:899–906.

de Kieviet JF, Zoetebier L, Van Elburg RM, Vermeulen RJ, Oosterlaan J. 2012. Brain development of very preterm and very low□birthweight children in childhood and adolescence: A meta□analysis. Dev Med Child Neurol. 54:313–323.

Dimitrova R, Pietsch M, Christiaens D, Ciarrusta J, Wolfers T, Batalle D, Hughes E, Hutter J, Cordero-Grande L, Price AN, Chew A, Falconer S, Vecchiato K, Steinweg JK, Carney O, Rutherford MA, Tournier J-D, Counsell SJ, Marquand AF, Rueckert D, Hajnal J V, McAlonan G, Edwards AD, O’Muircheartaigh J. 2020. Heterogeneity in Brain Microstructural Development Following Preterm Birth. Cereb Cortex. 30:4800–4810.

Duerden EG, Thompson DK. 2020. Can you see what I see? Assessing brain maturation and injury in preterm and term neonates. Brain. 143:383–386.

Edwards AD, Redshaw ME, Kennea N, Rivero-Arias O, Gonzales-Cinca N, Nongena P, Ederies M, Falconer S, Chew A, Omar O, Hardy P, Harvey ME, Eddama O, Hayward N, Wurie J, Azzopardi D, Rutherford MA, Counsell S. 2018. Effect of MRI on preterm infants and their families: a randomised trial with nested diagnostic and economic evaluation. Arch Dis Child Fetal Neonatal Ed. 103:F15–F21.

El Marroun H, Zou R, Leeuwenburg MF, Steegers EAP, Reiss IKM, Muetzel RL, Kushner SA, Tiemeier H. 2020. Association of Gestational Age at Birth With Brain Morphometry. JAMA Pediatr. 174:1149–1158.

Elovitz MA, Mrinalini C. 2004. Animal models of preterm birth. Trends Endocrinol Metab. 15:479–487.

George JM, Pagnozzi AM, Bora S, Boyd RN, Colditz PB, Rose SE, Ware RS, Pannek K, Bursle JE, Fripp J, Barlow K, Iyer K, Leishman SJ, Jendra RL. 2020. Prediction of childhood brain outcomes in infants born preterm using neonatal MRI and concurrent clinical biomarkers (PREBO-6): study protocol for a prospective cohort study. BMJ Open. 10:e036480.

Gui L, Loukas S, Lazeyras F, Hüppi PS, Meskaldji DE, Borradori Tolsa C. 2019. Longitudinal study of neonatal brain tissue volumes in preterm infants and their ability to predict neurodevelopmental outcome. Neuroimage. 185:728–741.

Holland D, Chang L, Ernst TM, Curran M, Buchthal SD, Alicata D, Skranes J, Johansen H, Hernandez A, Yamakawa R, Kuperman JM, Dale AM. 2014. Structural growth trajectories and rates of change in the first 3 months of infant brain development. JAMA Neurol. 71:1266–1274.

Hughes EJ, Winchman T, Padormo F, Teixeira R, Wurie J, Sharma M, Fox M, Hutter J, Cordero-grande L, Price AN, Allsop J, Bueno-conde J, Tusor N, Arichi T, Edwards AD, Rutherford MA, Counsell SJ, Hajnal J V. 2017. A Dedicated Neonatal Brain Imaging System. Magn Reson Med. 78:794–804.

Hüppi PS, Warfield S, Kikinis R, Barnes PD, Zientara GP, Jolesz FA, Tsuji MK, Volpe JJ. 1998. Quantitative magnetic resonance imaging of brain development in premature and mature newborns. Ann Neurol. 43:224–235.

Huttenlocher PR. 1990. Morphometric study of human cerebral cortex development. Neuropsychologia. 28:517–527.

Huttenlocher PR, Dabholkar AS. 1997. Regional differences in synaptogenesis in human cerebral cortex. J Comp Neurol. 387:167–178.

Inder TE, Warfield SK, Wang H, Hüppi PS, Volpe JJ. 2005. Abnormal cerebral structure is present at term in premature infants. Pediatrics. 115:286–294.

Kersbergen KJ, Makropoulos A, Aljabar P, Groenendaal F, de Vries LS, Counsell SJ, Benders MJNL. 2016. Longitudinal Regional Brain Development and Clinical Risk Factors in Extremely Preterm Infants. J Pediatr. 178:93–100.e6.

Knickmeyer RC, Gouttard S, Kang C, Evans D, Wilber K, Smith JK, Hamer RM, Lin W, Gerig G, Gilmore JH. 2008. A structural MRI study of human brain development from birth to 2 years. J Neurosci. 28:12176–12182.

Kostović I, Jovanov-Milošević N, Radoš M, Sedmak G, Benjak V, Kostović-Srzentić M, Vasung L, Čuljat M, Radoš M, Hüppi P, Judaš M. 2014. Perinatal and early postnatal reorganization of the subplate and related cellular compartments in the human cerebral wall as revealed by histological and MRI approaches. Brain Struct Funct. 219:231–253.

Kostovic I, Rakic P. 1990. Developmental history of the transient subplate zone in the visual and somatosensory cortex of the macaque monkey and human brain. J Comp Neurol. 297:441–470.

Lebenberg J, Mangin JF, Thirion B, Poupon C, Hertz-Pannier L, Leroy F, Adibpour P, Dehaene-Lambertz G, Dubois J. 2019. Mapping the asynchrony of cortical maturation in the infant brain: A MRI multi-parametric clustering approach. Neuroimage. 185:641–653.

Ligam P, Haynes RL, Folkerth RD, Liu L, Yang M, Volpe JJ, Kinney HC. 2009. Thalamic damage in periventricular leukomalacia: novel pathologic observations relevant to cognitive deficits in survivors of prematurity. Pediatr Res. 65:524–529.

Lin Y, Okumura A, Hayakawa F, Kato T, Kuno K, Watanabe K. 2001. Quantitative evaluation of thalami and basal ganglia in infants with periventricular leukomalacia. Dev Med Child Neurol. 43:481–485.

Makropoulos A, Aljabar P, Wright R, Hüning B, Merchant N, Arichi T, Tusor N, Hajnal J V., Edwards AD, Counsell SJ, Rueckert D. 2016. Regional growth and atlasing of the developing human brain. Neuroimage. 125:456–478.

Makropoulos A, Robinson EC, Schuh A, Wright R, Fitzgibbon S, Bozek J, Counsell SJ, Steinweg J, Vecchiato K, Passerat-Palmbach J, Lenz G, Mortari F, Tenev T, Duff EP, Bastiani M, Cordero-Grande L, Hughes E, Tusor N, Tournier JD, Hutter J, Price AN, Teixeira RPAG, Murgasova M, Victor S, Kelly C, Rutherford MA, Smith SM, Edwards AD, Hajnal J V., Jenkinson M, Rueckert D. 2018. The developing human connectome project: A minimal processing pipeline for neonatal cortical surface reconstruction. Neuroimage. 173:88–112.

Marquand AF, Kia SM, Zabihi M, Wolfers T, Buitelaar JK, Beckmann CF. 2019. Conceptualizing mental disorders as deviations from normative functioning. Mol Psychiatry. 24:1415–1424.

Marquand AF, Rezek I, Buitelaar J, Beckmann CF. 2016. Understanding Heterogeneity in Clinical Cohorts Using Normative Models: Beyond Case-Control Studies. Biol Psychiatry. 80:552–561.

Ment LR, Kesler S, Vohr B, Katz KH, Baumgartner H, Schneider KC, Delancy S, Silbereis J, Duncan CC, Constable RT, Makuch RW, Reiss AL. 2009. Longitudinal Brain Volume Changes in Preterm and Term Control Subjects During Late Childhood and Adolescence. Pediatrics. 123:503 LP–511.

Nosarti C. 2002. Adolescents who were born very preterm have decreased brain volumes. Brain. 125:1616–1623.

Nosarti C, Reichenberg A, Murray RM, Cnattingius S, Lambe MP, Yin L, MacCabe J, Rifkin L, Hultman CM. 2012. Preterm Birth and Psychiatric Disorders in Young Adult Life. Arch Gen Psychiatry. 69:610–617.

O’Muircheartaigh J, Robinson E, Pietsch M, Wolfers T, Aljabar P, Grande LC, Teixeira RPAG, Bozek J, Schuh A, Makropoulos A, Batalle D, Hutter J, Vecchiato K, Steinweg JK, Fitzgibbon S, Hughes E, Price A, Marquand A, Reuckert D, Rutherford M, Hajnal J, Counsell SJ, Edwards AD. 2020. Modelling brain development to detect white matter injury in term and preterm born neonates. Brain. 143:467–479.

Ou Y, Zöllei L, Retzepi K, Castro V, Bates S V., Pieper S, Andriole KP, Murphy SN, Gollub RL, Grant PE. 2017. Using clinically acquired MRI to construct age-specific ADC atlases: Quantifying spatiotemporal ADC changes from birth to 6-year old. Hum Brain Mapp. 38:3052–3068.

Petanjek Z, Judaš M, Kostović I, Uylings HBM. 2008. Lifespan Alterations of Basal Dendritic Trees of Pyramidal Neurons in the Human Prefrontal Cortex: A Layer-Specific Pattern. Cereb Cortex. 18:915–929.

Pierson CR, Folkerth RD, Billiards SS, Trachtenberg FL, Drinkwater ME, Volpe JJ, Kinney HC. 2007. Gray matter injury associated with periventricular leukomalacia in the premature infant. Acta Neuropathol. 114:619–631.

Sled JG, Nossin-Manor R. 2013. Quantitative MRI for studying neonatal brain development. Neuroradiology. 55:97–104.

Thompson DK, Matthews LG, Alexander B, Lee KJ, Kelly CE, Adamson CL, Hunt RW, Cheong JLY, Spencer-Smith M, Neil JJ, Seal ML, Inder TE, Doyle LW, Anderson PJ. 2020. Tracking regional brain growth up to age 13 in children born term and very preterm. Nat Commun. 11:696.

Towgood KJ, Meuwese JDI, Gilbert SJ, Turner MS, Burgess PW. 2009. Advantages of the multiple case series approach to the study of cognitive deficits in autism spectrum disorder. Neuropsychologia. 47:2981–2988.

Travis K, Ford K, Jacobs B. 2005. Regional Dendritic Variation in Neonatal Human Cortex: A Quantitative Golgi Study. Dev Neurosci. 27:277–287.

Tusor N, Benders MJ, Counsell SJ, Nongena P, Ederies MA, Falconer S, Chew A, Gonzalez-Cinca N, Hajnal J V., Gangadharan S, Chatzi V, Kersbergen KJ, Kennea N, Azzopardi D V., Edwards AD. 2017. Punctate White Matter Lesions Associated with Altered Brain Development and Adverse Motor Outcome in Preterm Infants. Sci Rep. 7:1–9.

Vasung L, Jovanov-Milošević N, Pletikos M, Mori S, Judaš M, Kostović I. 2011. Prominent periventricular fiber system related to ganglionic eminence and striatum in the human fetal cerebrum. Brain Struct Funct. 215:237–253.

Volpe JJ. 2009. Brain injury in premature infants: a complex amalgam of destructive and developmental disturbances. Lancet Neurol. 8:110–124.

Volpe JJ. 2019. Dysmaturation of Premature Brain: Importance, Cellular Mechanisms, and Potential Interventions. Pediatr Neurol. 95:42–66.

Wisnowski JL, Ceschin RC, Choi SY, Schmithorst VJ, Painter MJ, Nelson MD, Blüml S, Panigrahy A. 2015. Reduced thalamic volume in preterm infants is associated with abnormal white matter metabolism independent of injury. Neuroradiology. 57:515–525.

Wolfers T, Beckmann CF, Hoogman M, Buitelaar JK, Franke B, Marquand AF. 2019. Individual differences v. The average patient: Mapping the heterogeneity in ADHD using normative models. Psychol Med. 1–10.

Ziegler G, Ridgway GR, Dahnke R, Gaser C. 2014. Individualized Gaussian process-based prediction and detection of local and global gray matter abnormalities in elderly subjects. Neuroimage. 97:333–348.

