## Supplement for "Phenotyping the preterm brain: characterising individual deviations from normative volumetric development in two large infant cohorts"

**Supplementary Methods**:

**Exclusion Criteria for term-born infants:**

Exclusion criteria for the term-born infants included admission to neonatal intensive care unit (NICU) or significant intracranial abnormality detected on neonatal MRI scan including acute infarction and parenchymal haemorrhage. Term-born infants with punctate white matter lesions (PWMLs), mildly prominent ventricles or widening of the extra-axial CSF (within normative variation), small subependymal cysts, small caudothalamic cysts, small haemorrhages in the caudothalamic notch were not excluded from the study, as these are common findings on neonatal MRI. The term sample did not include infants from multiple pregnancies.

**Quality Control:**

T_2_-weighted images and tissue segmentation were examined to ensure no images severely affected by infant head motion or with poor segmentation were included in the present analysis. First, we assessed the success of the reconstruction and motion correction using a point scoring system, described in Makropoulos et al(Makropoulos et al. 2018). Upon visual examination, images of poor quality due to severe infant motion were assigned a score of 1, images with significant infant motion were ranked as 2, images of good quality with some negligible motion were rated as 3 and images with no visible effects of motion were ranked as 4 (*Supplementary* *Figure 1A*). Only images with scores 3 or 4 were included in the analysis. Furthermore, the visual examination ensured that the entire brain was captured within the field-of-view.

The quality of the tissue segmentation was examined in the same fashion. Infants were not included if the segmentation was unsuccessful (score 1) or poor (regional errors – score 2). As described in (Makropoulos et al. 2018), regional errors were mainly seen at the interface between the cerebellum and the inferior part of the occipital lobe, where part of the cortical ribbon was mislabelled as cerebellum caused by a poor image contrast. In other cases, deep WM, where the tissue intensity was very high, was mislabelled as ventricular volume (*Supplementary Figure 1B*). Only images with small localised segmentation errors (score 3) and those with no visible errors, hence good quality segmentation (score 4), were included in the analysis. No manual editing was used.

**Supplementary Figures:**

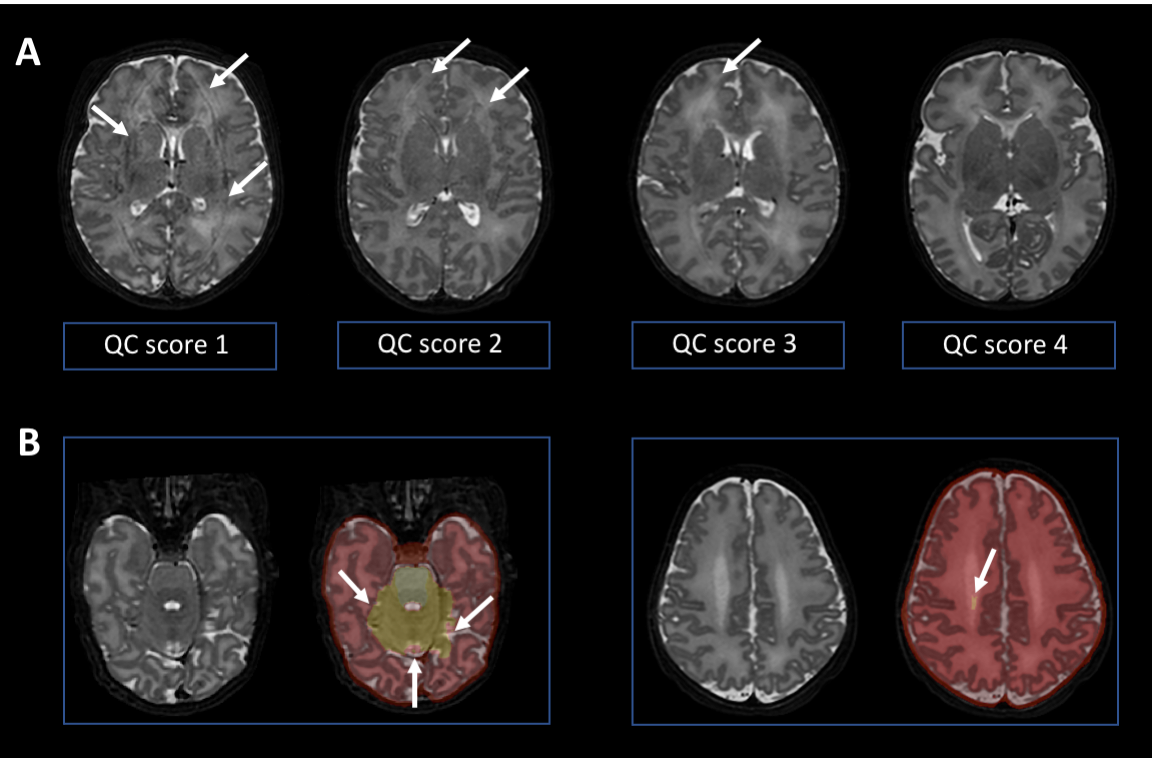

**Supplementary Figure 1.** **Quality control.** (A). Assessing the quality of motion correction: score 1 assigned to an image of poor quality; score 2 assigned to an image with significant infant motion; score 3 assigned to an image with some negligible motion; score 4 assigned to an image with good quality. (B) Assessing the quality of the segmentation: Examples of regional segmentation errors (score 2), where part of the cortical ribbon at the interface between the cerebellum and inferior temporal/occipital lobe is mislabelled as cerebellum due to the poor image contrast in this region or high intensity WM is mislabelled as ventricular volume. White arrows indicate motion artefacts or segmentation errors.

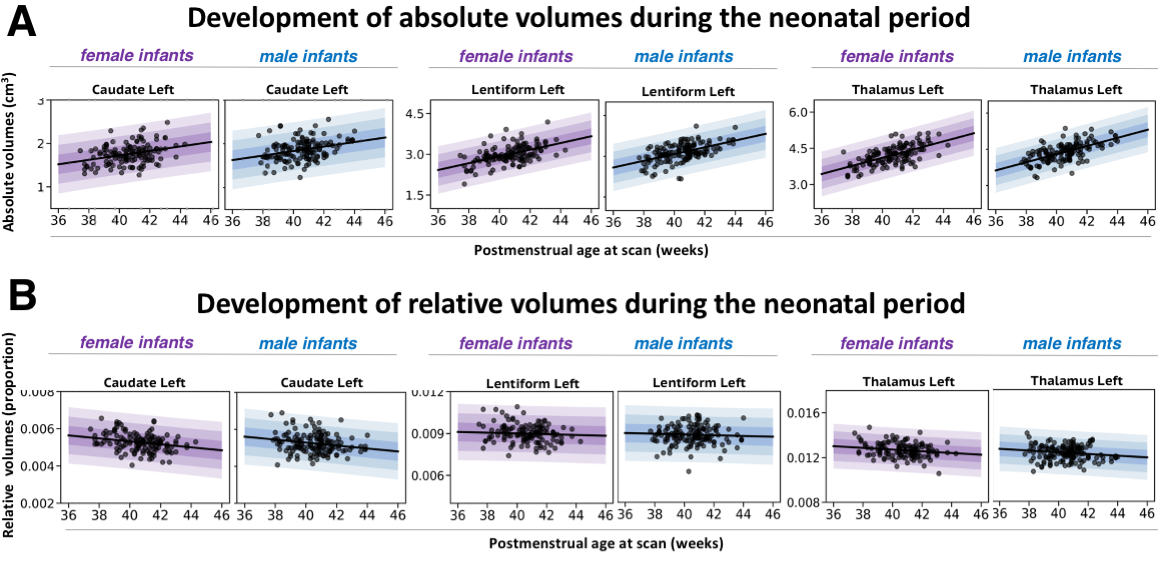

**Supplementary Figure 2.** **Normative modelling of the left subcortical volumes during the neonatal period.** Absolute volumes in cm^3^, relative volumes estimated as a proportion from TTV. The model means for both female and male infants are shown in purple and blue respectively together with ±1, ±2 and ±3 standard deviations from the model means.

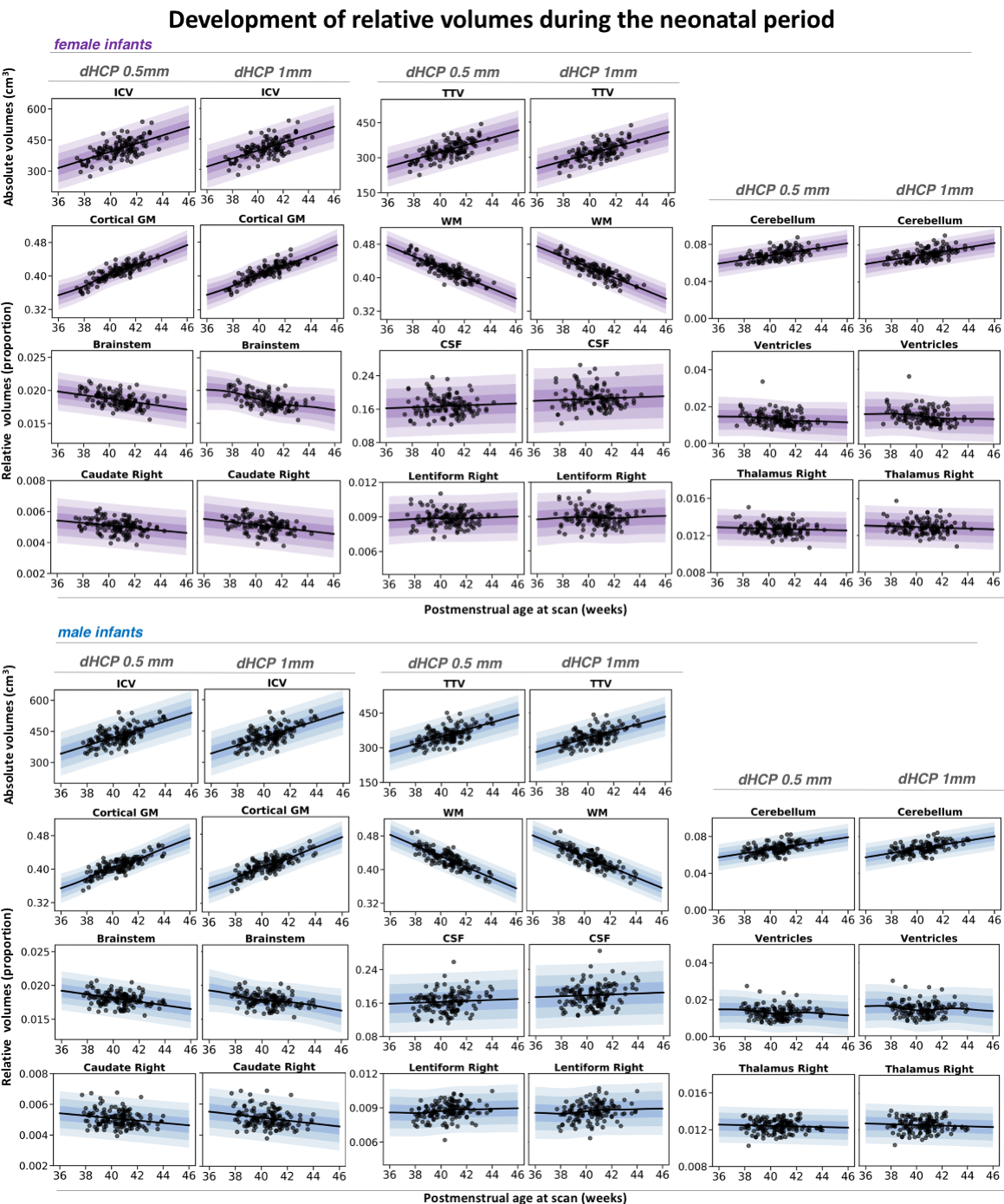

**Supplementary Figure 3.** **Normative modelling of volumetric development during the neonatal period using 0.5mm and 1mm resolution data.** The model means calculate for both voxel resolutions for both female and male term infants are shown in purple and blue respectively together with ±1, ±2 and ±3 standard deviations from the model means for relative volumes (tissue volumes represented as a proportion from TTV, ventricles from TBV and CSF from ICV).

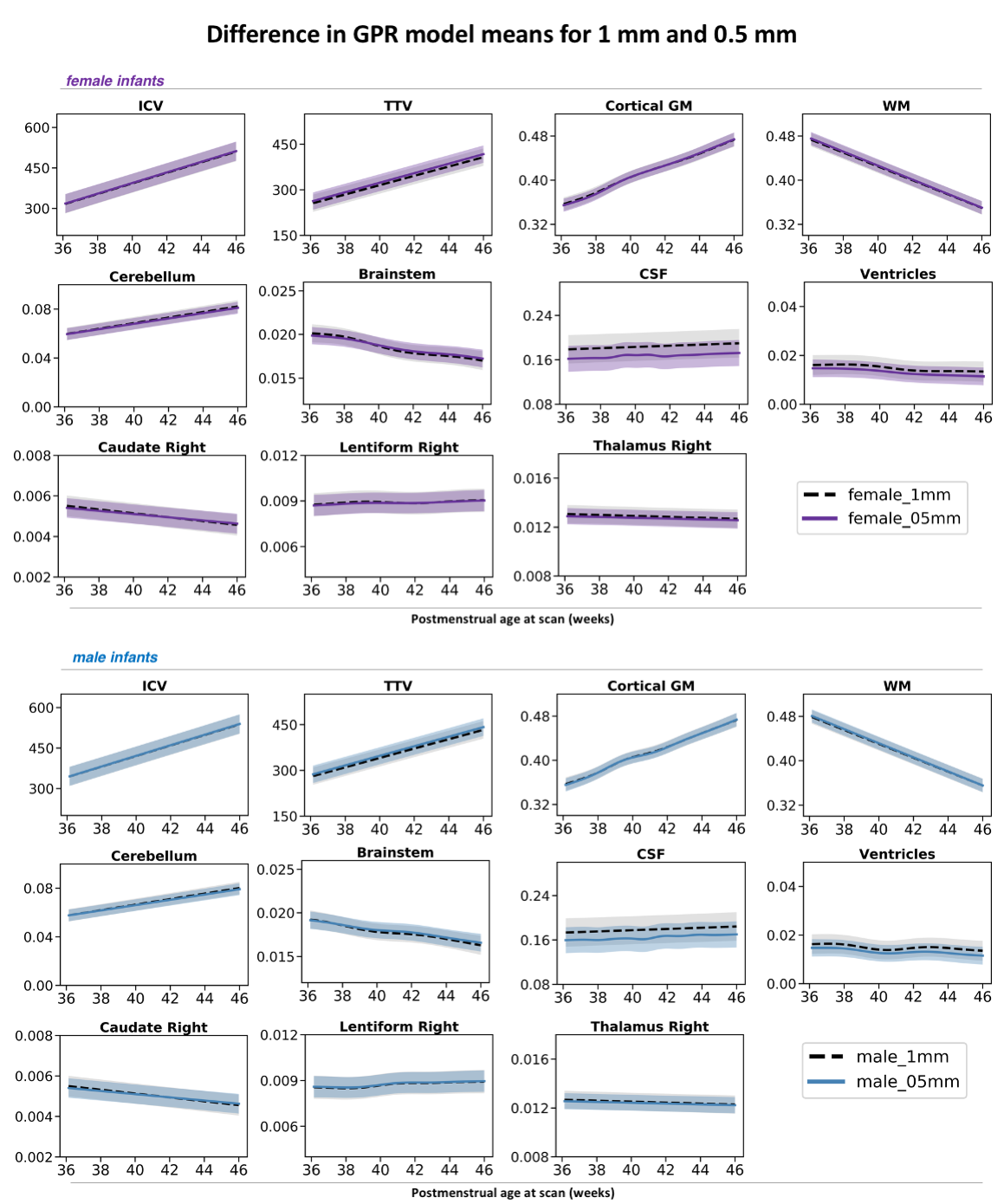

**Supplementary Figure 4. Effect of imaging voxel resolution on model means of neonatal volumetric development.** Model mean ±1 standard deviations are depicted separately for male and female infants in purple and blue respectively, and show a mean shift for fluid-filled structures (CSF and ventricles).

**Supplementary Tables:**

**Supplementary Table 1.** **Assessing the prediction of the normative models using train data under 5-fold cross-validation.** Mean absolute error (MAE) in units of standard deviation (sd) and Spearman ρ between observed and predicted volumes. In bold correlations significant after Bonferroni-Holm multiple comparison at p_corr_ < 0.05.

| **Brain region** | **Absolute volumes** | | **Relative volumes** | |
| --- | --- | --- | --- | --- |
|  | *MAE* | *Spearman ρ* | *MAE* | *Spearman ρ* |
| **ICV** | 0.59 | ***0.61*** | - | - |
| **TTV** | 0.58 | ***0.63*** | - | - |
| **CSF** | 0.73 | ***0.33*** | 0.78 | 0.10 |
| **cGM** | 0.49 | ***0.72*** | 0.43 | ***0.81*** |
| **WM** | 0.69 | ***0.44*** | 0.43 | ***0.83*** |
| **Ventricles** | 0.74 | 0 | 0.73 | 0.2 |
| **Cerebellum** | 0.48 | ***0.77*** | 0.65 | ***0.58*** |
| **Brainstem** | 0.63 | ***0.56*** | 0.69 | ***0.38*** |
| **Caudate Right** | 0.72 | ***0.37*** | 0.76 | ***0.22*** |
| **Caudate Left** | 0.72 | ***0.36*** | 0.76 | ***0.23*** |
| **Lentiform Right** | 0.61 | ***0.63*** | 0.76 | 0.10 |
| **Lentiform Left** | 0.64 | ***0.55*** | 0.77 | 0.10 |
| **Thalamus Right** | 0.61 | ***0.62*** | 0.77 | ***0.24*** |
| **Thalamus Left** | 0.59 | ***0.64*** | 0.77 | 0.19 |

**Supplementary Table 2. Correlation between deviations from the normative model and degree of prematurity (GA at birth).** Spearman rho, in bold correlations significant after Bonferroni-Holm multiple comparison at p_corr_<0.05.

| **Brain region** | **dHCP Term**  ***n = 274*** | **dHCP Preterm**  *n = 89* | **EPrime**  *n = 253* |
| --- | --- | --- | --- |
| **ICV** | -0.10 | 0.29 | 0.18 |
| **TTV** | -0.09 | ***0.45*** | ***0.27*** |
| **CSF** | 0.04 | ***-0.44*** | -0.10 |
| **cGM** | 0.02 | -0.17 | -0.02 |
| **WM** | -0.01 | 0.16 | 0.01 |
| **Ventricles** | -0.05 | -0.28 | ***-0.26*** |
| **Cerebellum** | 0.03 | -0.01 | 0.10 |
| **Brainstem** | 0.03 | 0.06 | 0 |
| **Caudate Right** | -0.03 | 0.19 | -0.08 |
| **Caudate Left** | -0.04 | 0.19 | -0.12 |
| **Lentiform Right** | 0.06 | -0.19 | 0.01 |
| **Lentiform Left** | -0.01 | -0.19 | 0.02 |
| **Thalamus Right** | 0.05 | -0.05 | -0.09 |
| **Thalamus Left** | 0.06 | -0.08 | -0.12 |

**Supplementary Table 3. Correlation between deviations from the normative model and birth weight Z-score.** Spearman rho, in bold correlations significant after Bonferroni-Holm multiple comparison at p_corr_ < 0.05.

| **Brain Regions** | **dHCP Term**  ***n = 274*** | **dHCP preterm**  *n = 89* | **EPrime**  *n = 245** |
| --- | --- | --- | --- |
| **ICV** | ***0.46*** | ***0.41*** | ***0.36*** |
| **TTV** | ***0.41*** | ***0.40*** | ***0.37*** |
| **CSF** | ***0.22*** | 0.12 | -0.10 |
| **cGM** | 0.07 | 0.09 | 0.16 |
| **WM** | 0.02 | 0 | -0.04 |
| **Ventricles** | 0.05 | -0.03 | 0.09 |
| **Cerebellum** | -0.11 | -0.08 | -0.12 |
| **Brainstem** | -0.14 | -0.13 | ***-0.26*** |
| **Caudate Right** | -0.03 | 0.01 | -0.06 |
| **Caudate Left** | 0.01 | 0.02 | -0.04 |
| **Lentiform Right** | -0.09 | 0.07 | -0.11 |
| **Lentiform Left** | -0.12 | 0.06 | -0.06 |
| **Thalamus Right** | -0.08 | -0.09 | ***-0.28*** |
| **Thalamus Left** | -0.11 | -0.12 | ***-0.25*** |

**Supplementary Table 4.** **Correlation between deviations from the normative model and oxygen support and nutrition.** Spearman rho, in bold correlations significant after Bonferroni-Holm multiple comparison at p_corr_ < 0.05. (Note TPN data were only available for EPrime)

| **Brain region** | **dHCP preterm**  *n = 89* | | **EPrime**  *n = 253* | | |
| --- | --- | --- | --- | --- | --- |
|  | *CPAP* | *ventilation* | *CPAP* | *ventilation* | *TPN* |
| **ICV** | ***-0.37*** | -0.30 | ***-0.21*** | ***-0.26*** | ***-0.35*** |
| **TTV** | ***-0.47*** | ***-0.44*** | ***-0.30*** | ***-0.38*** | ***-0.43*** |
| **CSF** | ***0.39*** | ***0.39*** | 0.11 | 0.16 | 0.07 |
| **cGM** | 0.17 | 0.20 | 0.04 | 0.08 | -0.06 |
| **WM** | -0.12 | -0.14 | -0.02 | -0.03 | 0.06 |
| **Ventricles** | ***0.35*** | ***0.42*** | ***0.22*** | ***0.22*** | ***0.25*** |
| **Cerebellum** | -0.02 | -0.01 | -0.05 | -0.13 | -0.09 |
| **Brainstem** | 0.02 | 0.10 | 0.06 | 0.05 | 0.17 |
| **Caudate Right** | -0.21 | -0.22 | -0.01 | 0 | 0.08 |
| **Caudate Left** | -0.23 | ***-0.34*** | 0.03 | 0.04 | 0.13 |
| **Lentiform Right** | -0.02 | 0.06 | -0.08 | -0.04 | -0.03 |
| **Lentiform Left** | -0.07 | -0.02 | -0.10 | -0.04 | -0.04 |
| **Thalamus Right** | 0.03 | 0.09 | 0.05 | 0.06 | 0.13 |
| **Thalamus Left** | 0 | 0.05 | 0.07 | 0.09 | 0.14 |

**Supplementary Table 5.** **Correlation between deviations from the normative model and behavioural follow-up.** Spearman rho, in bold, significant association after Bonferroni-Holm correction (at p_corr_ < 0.05).

| **Brain region** | **dHCP Term**  ***n = 222*** | | | **dHCP Preterm**  *n = 68* | | | **EPrime**  *n = 237* | | |
| --- | --- | --- | --- | --- | --- | --- | --- | --- | --- |
|  | **Mot** | **Lang** | **Cog** | **Mot** | **Lang** | **Cog** | **Mot** | **Lan** | **Cog** |
| **ICV** | -0.02 | 0.08 | 0.11 | 0.11 | 0.04 | 0.10 | 0.09 | 0.07 | 0.09 |
| **TTV** | 0 | 0.08 | 0.11 | 0.30 | 0.18 | 0.20 | 0.16 | 0.14 | ***0.20*** |
| **CSF** | -0.01 | 0.02 | 0 | -0.32 | -0.23 | -0.20 | -0.18 | ***-0.22*** | ***-0.25*** |
| **cGM** | 0.16 | 0.03 | 0.09 | -0.12 | 0.03 | -0.18 | -0.18 | ***-0.21*** | ***-0.20*** |
| **WM** | -0.09 | 0.07 | -0.03 | -0.08 | -0.18 | 0.02 | 0.11 | ***0.20*** | 0.14 |
| **Ventricles** | -0.20 | -0.04 | -0.07 | ***-0.40*** | ***-0.37*** | -0.32 | -0.16 | -0.08 | -0.12 |
| **Cerebellum** | -0.18 | -0.12 | -0.09 | 0.17 | 0.13 | 0.08 | 0.11 | 0.03 | 0.09 |
| **Brainstem** | -0.12 | -0.15 | -0.12 | 0.17 | 0.20 | 0.21 | 0.09 | 0.02 | 0.05 |
| **Caudate Right** | -0.08 | -0.07 | -0.03 | 0.11 | -0.06 | 0.06 | -0.05 | 0.06 | -0.07 |
| **Caudate Left** | -0.09 | -0.09 | -0.03 | 0.14 | -0.05 | 0.08 | -0.05 | 0.02 | -0.09 |
| **Lentiform Right** | -0.05 | -0.14 | -0.05 | 0.01 | -0.09 | -0.05 | 0.11 | 0.11 | 0.13 |
| **Lentiform Left** | -0.04 | -0.10 | 0 | 0.04 | -0.13 | -0.10 | 0.13 | 0.14 | 0.12 |
| **Thalamus Right** | 0.03 | -0.03 | -0.02 | 0 | 0.03 | 0.08 | 0.16 | 0.02 | 0.13 |
| **Thalamus Left** | 0.08 | 0.05 | -0.03 | 0.11 | 0.13 | 0.12 | 0.18 | -0.02 | 0.08 |

**References:**

Makropoulos A, Robinson EC, Schuh A, Wright R, Fitzgibbon S, Bozek J, Counsell SJ, Steinweg J, Vecchiato K, Passerat-Palmbach J, Lenz G, Mortari F, Tenev T, Duff EP, Bastiani M, Cordero-Grande L, Hughes E, Tusor N, Tournier JD, Hutter J, Price AN, Teixeira RPAG, Murgasova M, Victor S, Kelly C, Rutherford MA, Smith SM, Edwards AD, Hajnal J V., Jenkinson M, Rueckert D. 2018. The developing human connectome project: A minimal processing pipeline for neonatal cortical surface reconstruction. Neuroimage. 173:88–112.
